# Microstructural plasticity in the bilingual brain

**DOI:** 10.1101/657932

**Authors:** Daiyi Luo, Veronica P.Y. Kwok, Ping Li, Qing Liu, Wenlong Li, Yang Yang, Ke Zhou, Min Xu, Jia-Hong Gao, Li Hai Tan

## Abstract

The human brain has been uniquely equipped with the remarkable ability to acquire more than one language, as in bilingual individuals. Previous neuroimaging studies have indicated that learning a second language (L2) induced neuroplasticity at the macrostructural level. In this study, using the quantitative MRI (qMRI) combined with functional MRI (fMRI) techniques, we quantified the microstructural properties and tested whether second language learning modulates the microstructure in the bilingual brain. We found significant microstructural variations related to age of acquisition of second language in the left inferior frontal region and the left fusiform gyrus that are crucial for resolving lexical competition of bilinguals’ two languages. Early second language acquisition contributes to enhance cortical development at the microstructural level.

**Significant statement:** The ability to communicate in two languages is becoming more and more important in the increasingly global community. Does learning a second language (L2) affect the human brain development? At the macrostructural level, there has been neuroimaging evidence revealing neuroplasticity induced by the acquisition of L2. Here, we employed the quantitative MRI technique to investigate the microstructural variations related to L2 learning, and found that age of acquisition of L2, but not its proficiency, is associated with cortical proliferation. Early second language acquisition seems to enhance cortical development at the microstructural level.

## Introduction

One of the key characteristics of the bilingual brain is that when processing the target language, bilinguals need to successfully monitor and resolve lexical interference from the non-target language that competes for representation and selection (Crinion et al., 2006; Green et al., 2006; A. Hernandez et al., 2005; Kovelman et al., 2008; Price et al., 1999; Tan et al., 2011; Thierry & Wu, 2007; Xu et al., 2017). This has been argued to lead to cognitive advantages on executive tasks due to bilingualism (Bialystok et al., 2008; Bialystok et al., 2004; Birke Hansen et al., 2016; Colzato et al., 2008; Costa et al., 2008; Gold et al., 2013; Perani et al., 2017; Prior & MacWhinney, 2010). Past neuroimaging studies have demonstrated that learning a second language (L2) induced neuroplasticity at the macrostructural level, as indexed by gray matter density (Grogan et al., 2009; Mechelli et al., 2004), white matter integrity (Elmer et al., 2011; Hamalainen et al., 2017; Patricia K. Kuhl et al., 2016; Pliatsikas et al., 2015) and cortical thickness and volume (Klein et al., 2014; Li et al., 2014). Moreover, significant functional and structural imaging data points to the neural correlates of both L2 age of acquisition (AoA) and L2 proficiency. Early evidence suggests that childhood bilingualism may lead to distinct neural representations for L1 vs. L2, as compared with adulthood bilingualism (Kim et al., 1997). Later studies found out that proficiency, instead of AoA, may be the more important factor for determining the patterns of activation in L1 vs. L2 (Chee et al., 2001). It is unclear, however, whether effects due to AoA and proficiency can be separated or isolated, as age and proficiency are often confounded or conflated (Kim et al., 1997; Hernandez, 2013).

The neuroimaging measures used in previous studies, however, are qualitative because they are derived from uncalibrated T1-weighted images, which are sensitive to multiple features of tissue organization and microstructure (Mezer et al., 2013). To quantitatively evaluate microstructural properties in vivo, we employed the qMRI technique to compute the brain macromolecular tissue volume (MTV) and quantitative T1, which linearly contributes to iron and myelin concentrations (Stüber et al., 2014). As cell membranes and proteins account for the majority of brain macromolecules, MTV provides a valid approximation of myelin volume (Berman et al., 2018). Developmental decrease of T1 is thought to result from microstructural proliferation such as dendrite development, myelination and oligodendrocytes (Gomez et al., 2017). We used MTV and quantitative T1 to determine the microstructural variation in the brain tissue among bilinguals by manipulating AoA of second language (L2).

Fifty right-handed proficient Chinese-English bilinguals, including 25 early bilinguals (who learned L2 before age 6; 9 males and 16 females) and 25 late bilinguals (who learned English after age 9; 10 males and 15 females), participated in the fMRI (TR=2000 ms, TE=30 ms, 33 slices, flip angle=90°, voxel size= 3.5×3.5×4.2 mm) and qMRI experiments (4 spoiled gradient echo scans with different flip angles of 4°, 10°, 20° and 30°; 4 spin echo inversion recovery scans with different inversion times at 50, 400, 1200 and 2400 ms). We used whole-brain functional MRI runs to evaluate cortical responses to different categories of visual stimuli (Chinese characters, English words, faces and still checkerboards), while participants decided whether the two consecutively presented stimuli were the same. To examine the cortical responses to the bilinguals’ second language processing, we focused on the English words vs. English scrambled words contrast. In addition, all participants were administered a qualitative language experience and proficiency questionnaire (Marian et al., 2007) and English language proficiency tests.

To identify the relationships between bilingual processing and executive function, we asked participants to complete a series of cognitive tasks (Tan et al., 2005) which are known to be related to bilinguals’ language development (Li et al., 2014), including the nonverbal Raven IQ test, the similarities and comprehension subtests of the Wechsler Adult Intelligence Scale, component search, rapid automatized naming of numbers, the working memory test, phoneme counting, phoneme deletion, and the Stroop task (see Methods and Table 1). Forty-six participants (23 early bilinguals and 23 late bilinguals) completed the cognitive tasks. Four participants did not complete any of the mentioned cognitive tasks and were excluded from the brain-behavior correlation analysis. The Stroop task required the participants to name the ink color of Chinese and English color words (red, blue, green, yellow) in the congruent (i.e. the ink color was consistent with the meaning of the color words) and incongruent (i.e. the ink color was inconsistent with the meaning of the color words) conditions (MacLeod, 1991). In this task, participants needed to inhibit the competition of unrelated interfering information in the incongruent condition.

**Table 1.** Demographic characteristics and descriptive statistics for all participants.

|  | Early bilinguals (n=25) | Late bilinguals (n=25) |
| --- | --- | --- |
| Male/Female | 9/16 | 10/15 |
| Age | 21 y and 5 m (2 y) | 22 y and 7 m (2 y and 2m) |
| Age of L2 acquisition | 3 y and 8 m (1 y and 2 m) | 10 y and 3 m (1 y and 4 m) |
| Proficiency test (score) | 22.48 (6.20) | 18.40 (6.72) |
| Nonverbal IQ | 56.70 (3.14) | 54.87 (4.61) |
| WAIS similarities | 20.26 (2.22) | 21.04 (2.01) |
| WAIS comprehension | 20.65 (1.80) | 21.48 (2.06) |
| component search | 33.30 (6.80) | 29.22 (5.64) |
| RAN of numbers (s) | 25.96 (4.45) | 28.05 (4.92) |
| forward numeric working memory | 9.91 (1.20) | 8.87 (1.63) |
| backward numeric working memory | 7.78 (1.68) | 6.22 (1.73) |
| phoneme counting | 11.61 (4.95) | 9.64 (3.91) |
| phoneme deletion | 25.70 (3.57) | 23.73 (3.69) |
| ***Stroop test*** |  |  |
| *colored squares* |  |  |
| total response time (s) | 16.32 (3.10) | 17.03 (2.48) |
| error rate (%) | 0.43 (1.15) | 0.58 (1.29) |
| *congruent Chinese color words* |  |  |
| total response time (s) | 15.88 (3.90) | 15.80 (3.29) |
| error rate (%) | 0.43 (1.53) | 0.58 (1.64) |
| *incongruent Chinese color words* |  |  |
| total response time (s) | 24.41 (7.60) | 24.87 (4.69) |
| error rate (%) | 3.33 (3.89) | 2.03 (2.97) |
| *congruent English color words* |  |  |
| total response time (s) | 15.26 (3.97) | 16.89 (3.83) |
| error rate (%) | 0 (0) | 0.43 (1.53) |
| *incongruent English color words* |  |  |
| total response time (s) | 26.42 (5.72) | 30.41 (7.75) |
| error rate (%) | 1.30 (2.80) | 1.88 (2.81) |

Data are presented as mean, with standard deviations in parentheses.

## Materials and Methods

### Ethics statement

The ethics board of the Shenzhen Institute of Neuroscience approved the study. All subjects provided written informed consent before the experiment.

### Subjects

A total of 50 proficient bilingual subjects who were Chinese native speakers and learned English as a second language (L2) participated in this study, including 25 early bilinguals who learned English before age of 6 (mean age 21 y and 5 m with standard deviation at 2 y, 9 males and 16 females) and 25 late bilinguals who learned English after age of 9 (mean age 22 y and 7 m with standard deviation at 2 y and 2 m, 10 males and 15 females). All subjects were right-handed, with normal or corrected-to-normal vision, physically healthy and neurologically typical young adults, with no alcohol or substance abuse or dependence.

### Language experience and proficiency

To assess the language experience of the participants, subjects completed the qualitative Language Experience and Proficiency Questionnaire (Marian et al., 2007). We used the listening and reading sub-sections of the International English Language Testing System (IELTS) as the English proficiency test to evaluate participants’ proficiency of L2. The maximum score on the proficiency test is 34.

### Cognitive tests

All cognitive tasks were administered individually. Among 50 recruited subjects, four subjects did not complete any of the tasks, and a total of 46 subjects (23 early bilinguals and 23 late bilinguals) participated in the cognitive tasks. One late bilingual completed all tasks except phoneme counting and phoneme deletion. and 45 subjects (23 early bilinguals and 22 late bilinguals) who completed all the tasks.

#### Nonverbal Raven IQ test

The standard Chinese version of Raven’s Standard Progressive Matrices (Raven, 1996) was used to measure the nonverbal intelligence of subjects. The maximum score is 60.

#### Subtests of the WAIS

Similarities: This test consisted of 13 pairs of Chinese words for object, direction, or behavior (e.g., “斧头(axe)” and “锯子(saw)”). The subjects were asked to tell the similarities among the pair of words. The experimenter rated the answers from 0 to 3. The test ended when the subject got 0 point for 4 consecutive times.

Comprehension: This test consisted of 14 questions concerning social values, social customs and reasons for certain phenomena (e.g., “Why do we need traffic police in cities?”). The subjects were asked to answer the questions. The experimenter rated the answers from 0 to 2. The test ended when the subject got 0 point for 4 consecutive times.

#### Component search

150 Chinese characters among which some characters contain the component “木” or “又” were presented to the subjects (e.g., the character “集” contains the component “木”; the character “叔” contains the component “又”). The subjects were asked to find out the characters containing the component “木” or “又” and circle it as fast and precisely as possible in 80 s. The number of correctly circled characters was recorded for each subject.

#### Rapid automatized naming of numbers

Subjects were shown 100 single digits and required to name the digits in Chinese as fast as possible in sequence, from left to right and top to bottom. The completion time of each subject was recorded with a stopwatch.

#### Numeric working memory test

##### Forward digit-span task

The experimenter first read out a series of random digits at a speed of about one word per second. Subjects were then asked to recall the items accurately in the same order as they heard. The series of number was increasingly longer in each trial until the second time when the subject failed to recall the items correctly. The longest numbers of sequential digits were recorded for all subjects.

##### Backward digit-span task

The same as forward digit-span but the subjects need to recall the sequential digits in the reverse order of presentation.

#### Phoneme counting task

The experimenter read out 30 English words and, for each word, the subjects were required to answer the number of phonemes the word contains (e.g., the word “cake” contains 3 phonemes, /k/, /ei/ and /k/). The numbers of correct answers for all subjects were recorded.

#### Phoneme deletion task

The experimenter read out 30 English words and, for each word, told the subject to delete a specific phoneme and then enounce it (e.g., the phoneme /p/ was required to be deleted from the word “pear”, the correct answer is /eə/). The numbers of correct answers for all subjects were recorded.

#### The Stroop tasks

The subjects were required to name the ink color of each item as fast and precisely as possible. Each item was printed in red, blue, green or yellow. The Stroop task contained 5 types of stimuli, including neutral condition, Chinese congruent condition, Chinese incongruent condition, English congruent condition and English incongruent condition. In the neutral condition, colored squares were presented. Chinese and English color words (red, blue, green, yellow) were printed in the four colors, generating the congruent conditions, where the ink color was consistent with the meaning of the word, and the incongruent conditions, where the ink color was inconsistent with the meaning of the word. Each stimulus contained 30 trials. For all subjects, number of errors and total naming time in each condition were recorded (MacLeod, 1991).

### Data acquisition

#### Quantitative MRI

MRI experiments were performed on a 3 T Discovery MR750 system (General Electric Healthcare, Milwaukee, WI, USA) with an 8-channel head coil. Quantitative MRI measurements were obtained from the protocols in (Mezer et al., 2013). The quantitative MTV and T1 values were measured from spoiled gradient echo (SPGE) images with different flip angles of 4°, 10°, 20° and 30° (TR=14 ms, TE=2 ms) using 1×1 mm^2^ in-plane resolution with a slice thickness of 1 mm. For T1 calibration, four spin echo inversion recovery (SEIR) images were scanned, done with an echo planar imaging (EPI) read-out, a slab inversion pulse and spectral spatial fat suppression. Four SEIR images had different inversion times at 50, 400, 1200 and 2400 ms (TE=43 ms, TR=3.0 s) with a 2×2 mm^2^ in-plane resolution and a 4 mm slice thickness.

#### Functional MRI

Data were collected with the same scanner described above. Functional data were acquired with a T2*-weighted gradient-echo EPI sequence (TR=2000 ms, TE=30 ms, 33 slices, flip angle=90°, voxel size= 3.5×3.5×4.2 mm, FOV = 224 mm, interleaved slice order). Visual stimuli were presented through a projector onto a translucent screen and subjects viewed the screen through a mirror attached to the head coil.

#### FMRI category localizer experiment

To evaluate subjects’ cortical responses to different categories of visual stimuli, they viewed stimuli from 4 categories (Chinese characters, English words, faces and still checkerboards), while 2 categories contained two subcategories (Chinese characters: real characters and scrambled characters; English words: real words and scrambled words). Stimuli from each subcategory were presented in 20-s blocks, each subcategory block was alternated by a 22s-rest block. In each block, participants were instructed to judge whether two consecutively presented stimuli were the same. Each experimental block began with a 2-s instruction, followed by twelve 1.5-s trials. On each trial, the first image of a character/word/face/checkerboard was presented for 200 ms, followed by the presentation of a 200 ms fixation cross; the second image of a character/word/face/checkerboard was displayed for 500 ms. After that, another fixation cross was displayed for 600 ms for subjects to make a button press response. Presentation order of stimulus types was counterbalanced. Each run lasted 8 min 36 s, and each subject completed 3 runs. Since we here focused on second language processing, our data analysis was based only on the contrast of English words and English scrambled words.

### Data analysis

#### QMRI data analysis

Both SPGE images and SEIR images were processed using the mrQ software package (https://github.com/mezera/mrQ) to generate macromolecular tissue volume (MTV) map and quantitative T1 map for each subject. Unbiased T1 maps and proton density maps were estimated (Fram et al., 1987) by combining SPGE images and low-resolution unbiased T1 maps derived from SEIR images (Barral et al., 2010). MTV maps quantify the non-water volume in each voxel and they were estimated from proton density maps while cerebrospinal fluid was approximated to water. T1-weighted images, which were spatially matched with MTV maps and had excellent gray/white matter contrast, were processed using Freesurfer 6.0 recon-all procedure (Reuter et al., 2012).

#### FMRI data analysis

The fMRI data analysis was performed in MATLAB using SPM12 software package (http://www.fil.ion.ucl.ac.uk/spm). Functional data were corrected for slice-timing and realigned to the mean of the functional scans to remove movement artifact. Any participants who moved more than 2 mm within a scan were excluded from data analysis. The motion-corrected scans were then spatially normalized to the MNI standard space using individual high-resolution T1 anatomical images and resampled into 2×2×2 mm^3^ cubic voxels. The normalized images were then spatially smoothed with a 6mm isotropic Gaussian kernel. Individual activation maps were generated by using the general linear model (GLM) by convolving the experimental design with the hemodynamic response function (HRF), high-pass filtered at 128s, and the six head motion parameters were included as nuisance regressors. We used the GLM to generate statistical maps of contrasts between different conditions at the individual and group levels. Whole brain activation of contrast of interest (English words > English scrambled words) was computed by using a one-sample t test (*p* < 0.05, familywise error (FWE) corrected; extent threshold = 10). Brain regions and coordinates were reported in MNI space.

#### Definition of region of interest

We mainly focused on the left anterior inferior frontal cortex, left middle fusiform gyrus and anterior cingulate cortex as these regions were strongly activated in the functional task and were associated with language conflict monitoring in bilingual speakers. In these three mentioned regions, brain activity was found in statistical contrast of English words vs. English scrambled words at group level. The peak MNI coordinates in our task located closely to those reported in the literature that are important for resolving lexical competition (left anterior inferior frontal: peak (MNI: −48, 28, 12) vs. published (MNI: −44, 28, 8; BA 45/9) (Rodriguez et al., 2002); left middle fusiform: peak (MNI: −48, −44, −12) vs. published (MNI: −46, −57, −11; BA37) (Tan et al., 2011); anterior cingulate: peak (MNI: −4, −8, 40) vs. published (MNI: 5, 15, 40) (Abutalebi et al., 2012). The functional peak maxima were in the same anatomical location as the published coordinates, so we identified these three strongly activated regions of interest (ROIs) for qMRI analyses. For each ROI, the MNI coordinate was projected to fsaverage surface using the RF-ANTs MNI152-to-fsaverage mappings provided in (Wu et al., 2018) (https://github.com/ThomasYeoLab/CBIG/tree/master/stable_projects/registration/Wu2017_RegistrationFusion) and the nearest 10 vertices were selected to create a 2D surface label. The label was then dilated 8 times using Freesurfer’s mris_label_calc and converted to each subject’s individual cortical surface using mri_label2label. For each subject, by using mri_label2vol, the individual surface label was converted into a volumetric binary ROI mask (left anterior inferior frontal region averaged 1023 voxels, left fusiform averaged 1750 voxels, left anterior cingulate averaged 684 voxels) in the native MTV space, sampling a 3-mm thick ribbon below the gray-white matter boundary. The individual ROI masks were applied to the corresponding MTV maps and T1 maps for all subjects. Average MTV and T1 value across voxels within ROI for each subject was computed.

### Statistical analysis

Using one-way analysis of variance (ANOVA) and two-tailed t-tests in IBM SPSS, we assessed the group difference (early bilinguals vs. late bilinguals) in MTV and T1 values in each ROI. Group difference was considered to be significant at *p* < 0.05. To examine if MTV and T1 values within ROIs were correlated with L2 proficiency or age of L2 acquisition, we computed the Pearson correlation coefficient (*r*) and *p* value between them. When we analyzed the qMRI maps, data points that were 2 times of interquartile range (IQR) away from the group median were classified as outliers and were excluded from further analysis. Two late bilinguals were excluded from the left anterior inferior frontal region analysis, two early bilinguals and one late bilingual were excluded from the left middle fusiform analysis, and one late bilingual was excluded from the left cingulate analysis. Since qMRI measures were correlated with age of L2 acquisition in the left frontal and left fusiform region, we further computed the Pearson correlation coefficient (*r*) and *p* value between qMRI measures and performances in cognitive tasks. As for the Stroop task, Pearson partial correlations between qMRI measures and total naming time were calculated, with number of errors, age and proficiency as covariates. The alpha level was set at *p* < 0.017 (= 0.05/3 dependent measures) using the Bonferroni correction to correct for multiple comparisons.

## Results

Using fMRI, we identified three regions which were strongly activated in the functional task in the whole brain-based analysis (survived at p < 0.05 familywise error (FWE) correction threshold; see Table 2): the left anterior inferior frontal cortex (MNI: −48, 28, 12), the left fusiform (MNI: −48, −44, −12) and the anterior cingulate (MNI: −4, −8, 40) regions. The activation peak coordinate in each region was located closely to the coordinate reported in the existing literature on language interference in bilingualism: the left anterior inferior frontal region (MNI: −44, 28, 8; BA 45/9) (Rodriguez-Fornells et al., 2002), left middle fusiform region (MNI: −46, −57, −11; BA 37) (Tan et al., 2011) and anterior cingulate region (MNI: 5, 15, 44) (J. Abutalebi et al., 2012). These regions not only are the functional neural correlates of bilingualism but also have their neuroanatomical substrates (Li et al., 2014). We thus defined these 3 regions of interest (ROI) for qMRI data analyses, as shown in **Figure 1**. MTV and quantitative T1 maps were calculated for each subject (Mezer et al., 2013) using qMRI data. Average MTV and T1 values within the ROIs were computed to measure the microstructural proliferation of the bilingual participants. Then we evaluated the Pearson correlation coefficient between qMRI measures and AoA.

**Table 2.**
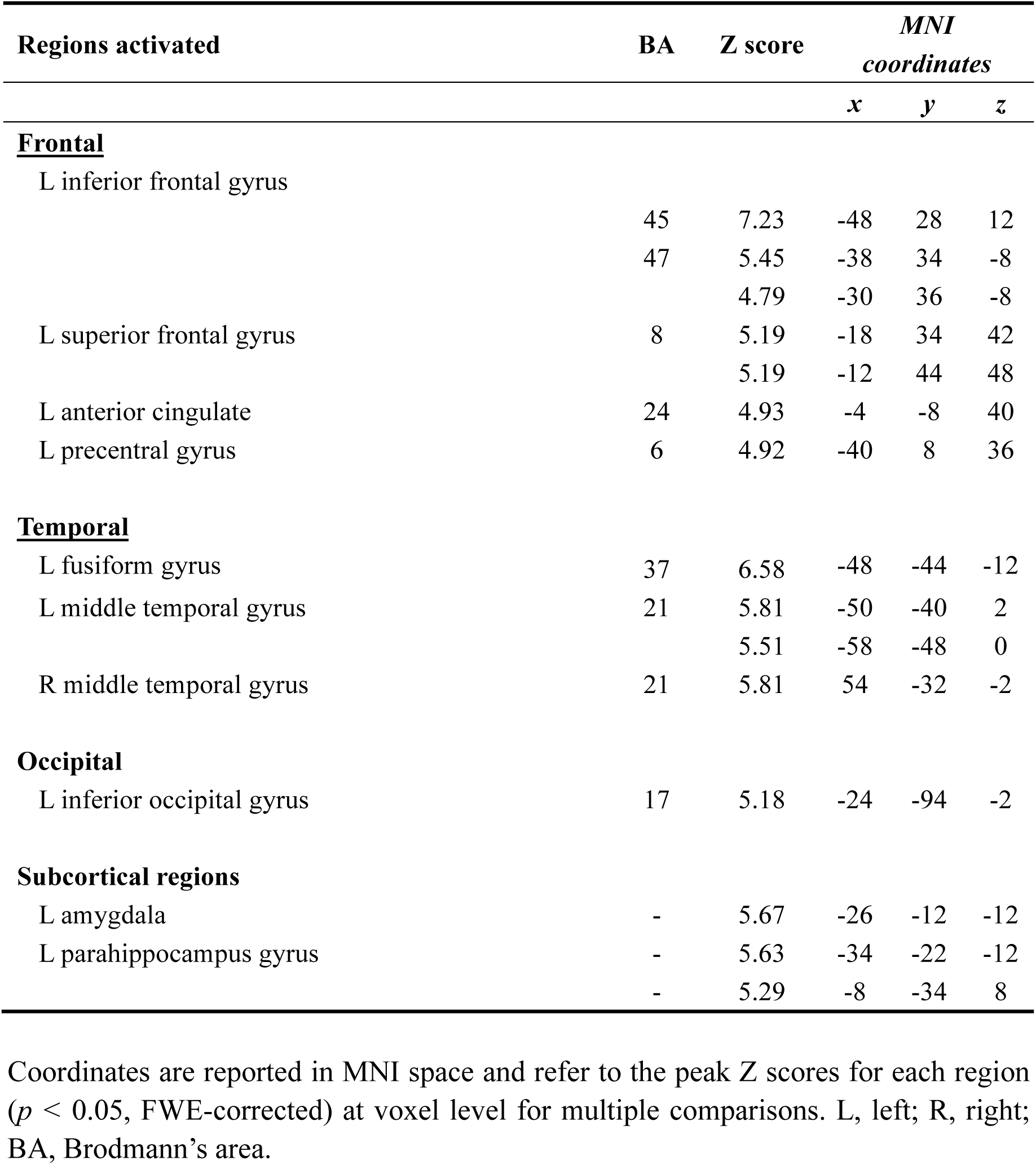
Coordinates of activation peaks: English real words condition minus English scrambled words condition.

| Regions activated | BA | Z score | *MNI*<br>*coordinates* |  |  |
| --- | --- | --- | --- | --- | --- |
|  |  |  | *x* | *y* | *z* |
| **<u>Frontal</u>** |  |  |  |  |  |
| L inferior frontal gyrus | 45 | 7.23 | -48 | 28 | 12 |
|  | 47 | 5.45 | -38 | 34 | -8 |
|  |  | 4.79 | -30 | 36 | -8 |
| L superior frontal gyrus | 8 | 5.19 | -18 | 34 | 42 |
|  |  | 5.19 | -12 | 44 | 48 |
| L anterior cingulate | 24 | 4.93 | -4 | -8 | 40 |
| L precentral gyrus | 6 | 4.92 | -40 | 8 | 36 |
| **<u>Temporal</u>** |  |  |  |  |  |
| L fusiform gyrus | 37 | 6.58 | -48 | -44 | -12 |
| L middle temporal gyrus | 21 | 5.81 | -50 | -40 | 2 |
|  |  | 5.51 | -58 | -48 | 0 |
| R middle temporal gyrus | 21 | 5.81 | 54 | -32 | -2 |
| **<u>Occipital</u>** |  |  |  |  |  |
| L inferior occipital gyrus | 17 | 5.18 | -24 | -94 | -2 |
| **<u>Subcortical regions</u>** |  |  |  |  |  |
| L amygdala | - | 5.67 | -26 | -12 | -12 |
| L parahippocampus gyrus | - | 5.63 | -34 | -22 | -12 |
|  | - | 5.29 | -8 | -34 | 8 |

Coordinates are reported in MNI space and refer to the peak Z scores for each region ( $p < 0.05$ , FWE-corrected) at voxel level for multiple comparisons. L, left; R, right; BA, Brodmann's area.

**Figure 1.**
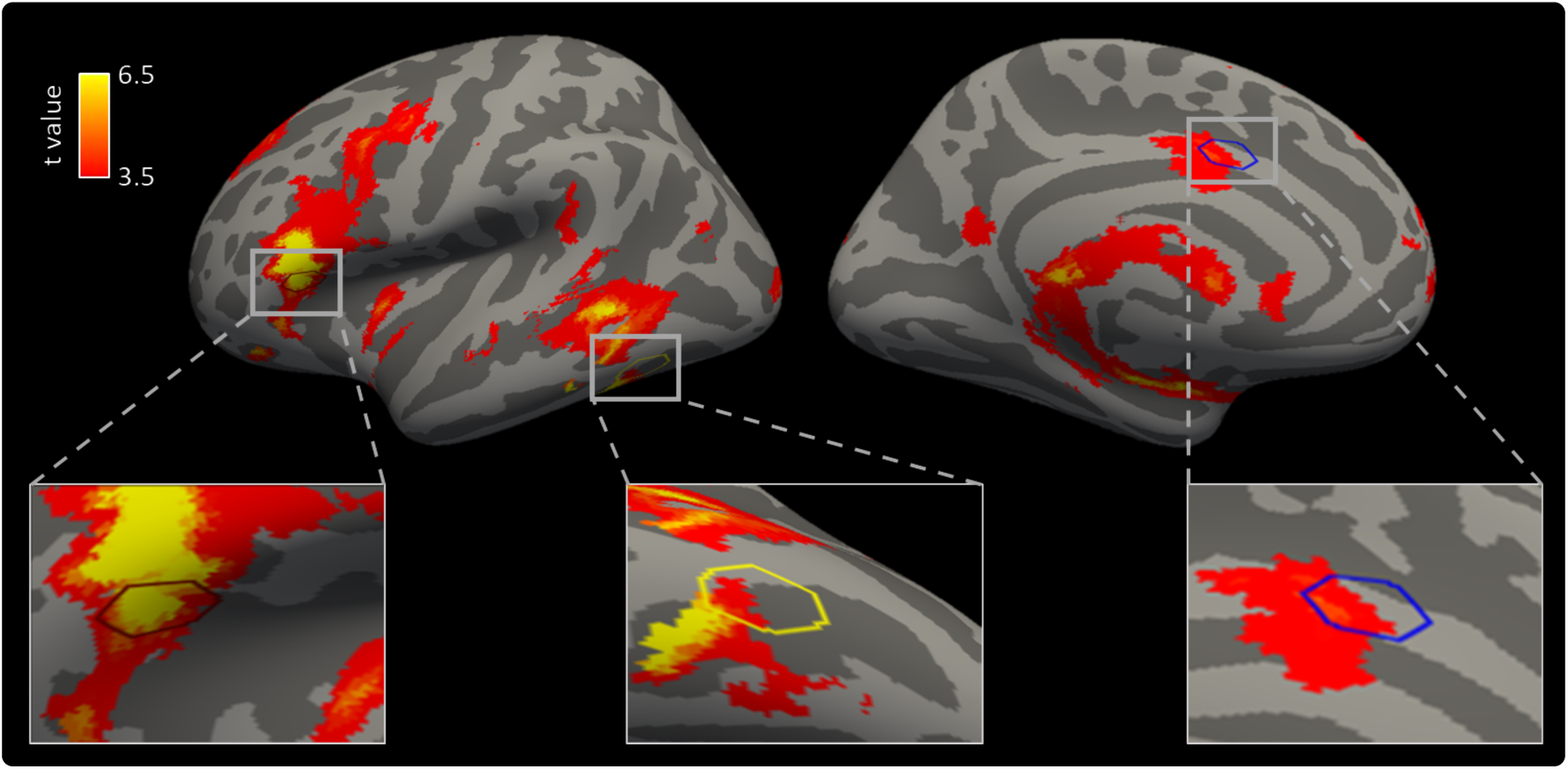
Averaged BOLD activation maps and regions of interest (ROI). Three ROIs were placed in left anterior inferior frontal region (the red hexagon), in left middle fusiform gyrus (the yellow hexagon) and in anterior cingulate region (the blue hexagon).

Significant microstructural variations related to AoA were identified in the left anterior inferior frontal region and left middle fusiform region. In the left anterior inferior frontal region, mean MTV was significantly higher (**Figure 2A**, *t*(46) = 2.967, *p* = 0.005) and mean T1 was significantly lower (**Figure 2D**, *t*(46) = −2.827, *p* = 0.007) in early bilinguals than in late bilinguals. We also found a reliable negative correlation between MTV and AoA (**Figure 2B**, *r* = −0.398, *p* = 0.006) and a positive correlation between T1 and AoA (**Figure 2E**, *r* = 0.409, *p* = 0.004). The analyses were repeated with language proficiency as a covariate, and the partial correlation remained significant between MTV and AoA (*r* = −0.396, *p* = 0.006), as well as between T1 and AoA (*r* = 0.375, *p* = 0.009).

**Figure 2.**
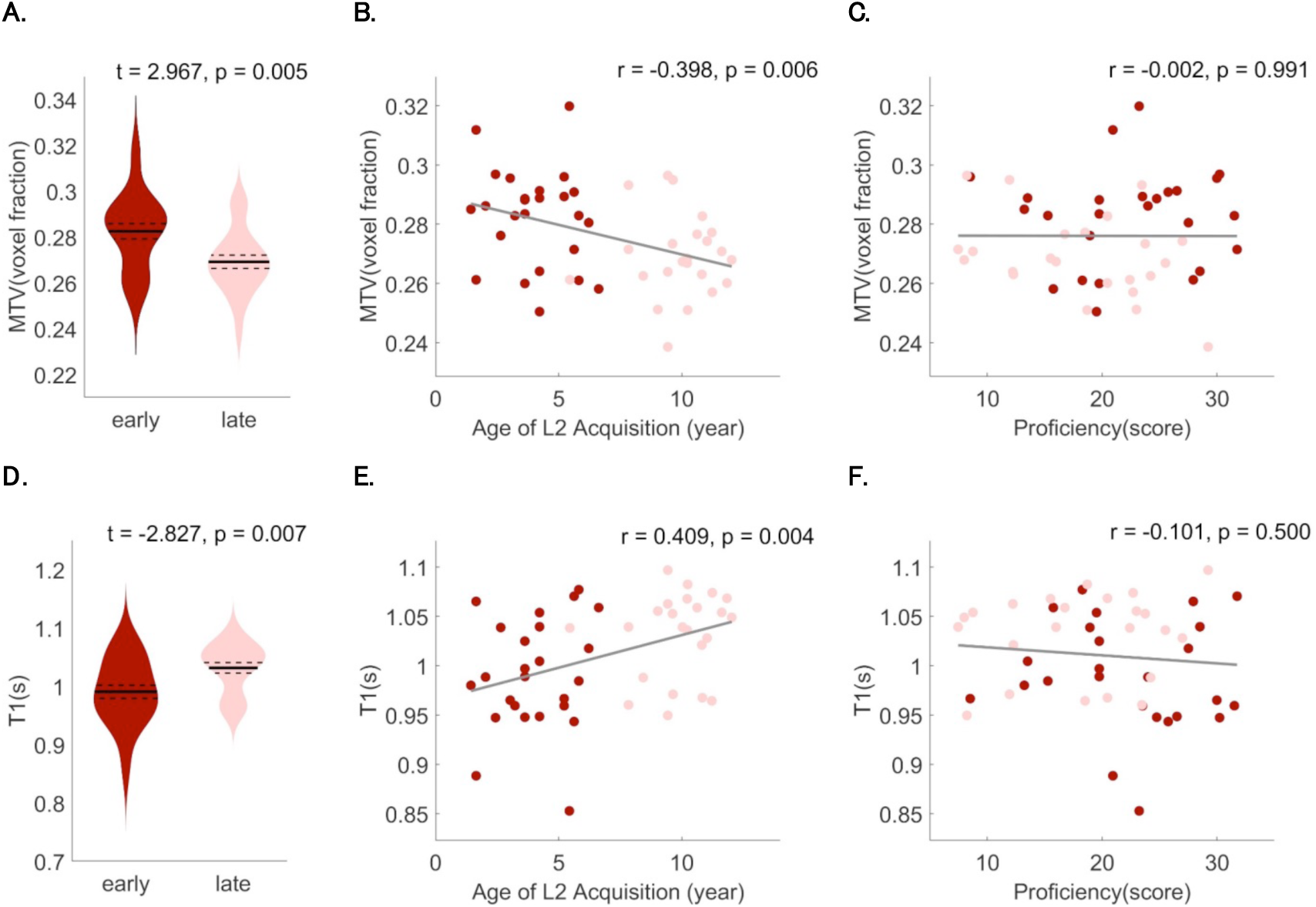
qMRI measures in ROI in left anterior inferior frontal region. (A) Average MTV in early and late bilingual groups. Violin plot shows the average MTV values across participants in the two groups. The width of the plot represents the participant distribution density within the group, the solid lines represent the group mean, the dotted lines represent the group standard error. (B) Correlation between MTV and age of L2 acquisition. Red dots represent early bilinguals (n=25) and pink dots represent late bilinguals (n=23). (C) Correlation between MTV and proficiency. (D) Average T1 in early and late bilingual groups. Violin plot shows the average T1 values across participants in the two groups. The width of the plot represents the participant distribution density within the group, the solid lines represent the group mean, the dotted lines represent the group standard error. (E) Correlation between T1 relaxation and age of L2 acquisition. (F) Correlation between T1 relaxation and proficiency.

In the left middle fusiform region, significant higher MTV (**Figure 3A**, *t*(45) = 2.932, *p* = 0.005) and lower T1 values (**Figure 3D**, *t*(45) = −2.155, *p* = 0.037) were observed in early bilinguals compared with late bilinguals. Correlation analyses indicated that MTV was negatively correlated with AoA (**Figure 3B**, *r* = −0.401, *p* = 0.006) while T1 was positively correlated with AoA (**Figure 3E**, *r* = 0.344, *p* = 0.019) in the left middle fusiform region. When language proficiency was added as a covariate, partial correlation also showed significant negative correlation between MTV and AoA (*r* = −0.419, *p* = 0.004) and positive correlation between T1 and AoA (*r* = 0.338, *p* = 0.021).

**Figure 3.**
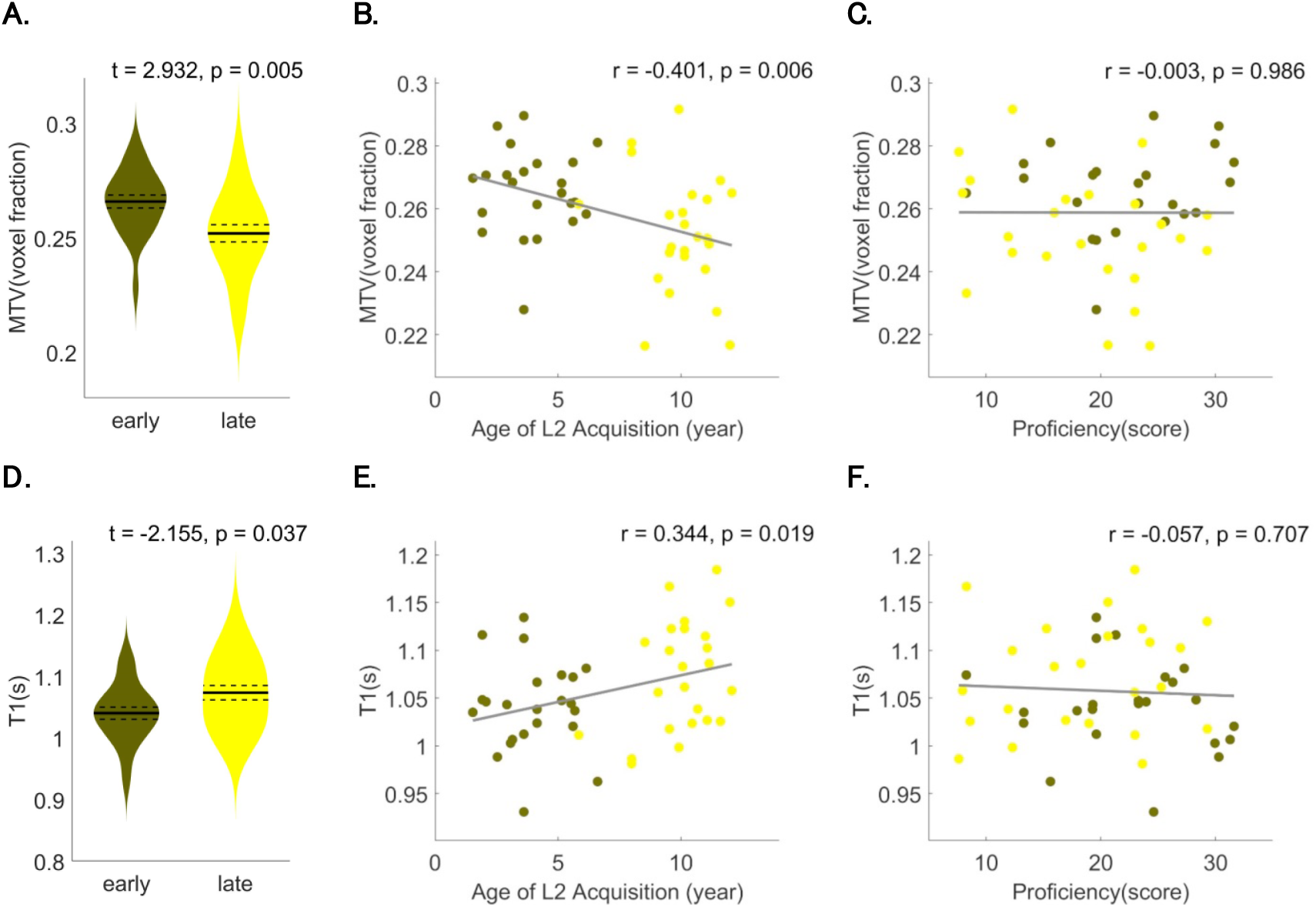
qMRI measures in ROI in left middle fusiform region. (A) Average MTV in early and late bilingual groups. Violin plot shows the average MTV values across participants in the two groups. The width of the plot represents the participant distribution density within the group, the solid lines represent the group mean, the dotted lines represent the group standard error. (B) Correlation between MTV and age of L2 acquisition. Olive dots represent early bilinguals (n=23) and yellow dots represent late bilinguals (n=24). (C) Correlation between MTV and proficiency. (D) Average T1 in early and late bilingual groups. Violin plot shows the average T1 values across participants in the two groups. The width of the plot represents the participant distribution density within the group, the solid lines represent the group mean, the dotted lines represent the group standard error. (E) Correlation between T1 relaxation and age of L2 acquisition. (F) Correlation between T1 relaxation and proficiency.

In the left anterior cingulate cortex, MTV showed a negative correlation trend with AoA (*p* = 0.114) while T1 showed a positive trend (*p* = 0.182). However, no significant difference in qMRI measures between early and late bilinguals was seen in this region (**Figure 4**).

**Figure 4.**
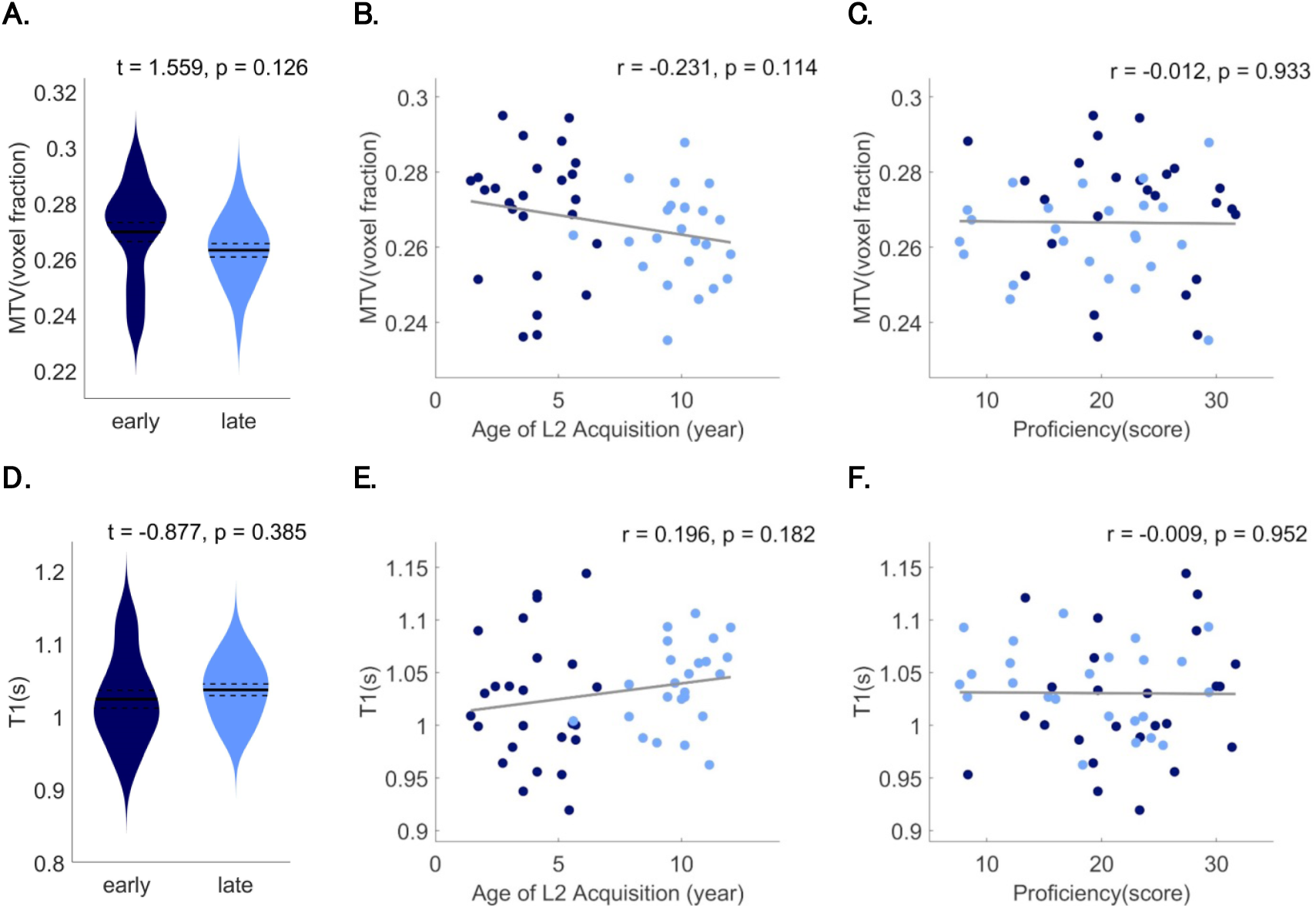
qMRI measures in the left anterior cingulate region. (A) Average MTV in early and late bilingual groups (*t*(47) = 1.559, *p* = 0.126). Violin plot shows the average MTV values across participants in the two groups. The width of the plot represents the participant distribution density within the group, the solid lines represent the group mean, the dotted lines represent the group standard error. (B) Correlation between MTV and age of L2 acquisition. Indigo dots represent early bilinguals (n=25) and blue dots represent late bilinguals (n=24). (C) Correlation between MTV and proficiency. (D) Average T1 relaxation time in early and late bilingual groups (*t*(47) = −0.877, *p* = 0.385). Violin plot shows the average T1 values across participants in the two groups. The width of the plot represents the participant distribution density within the group, the solid lines represent the group mean, the dotted lines represent the group standard error. (E) Correlation between T1 relaxation and age of L2 acquisition. (F) Correlation between T1 relaxation and proficiency.

Given these results, we further examined the relationships between qMRI measures and performances in cognitive tasks in the left frontal and left fusiform regions only. In the Stroop task, we examined the partial correlations between response time and qMRI measures with number of errors, actual age, and proficiency as covariates. Crucially, for the incongruent English color-words, there was a significant negative correlation between response time and MTV (**Figure 5A**, *r* = −0.344, *p* = 0.028) and a positive correlation between response time and T1 (**Figure 5B**, *r* = 0.336, *p* = 0.032) in the left frontal region. No significant correlations were found for other tasks. In the left middle fusiform, no significant correlations were identified between qMRI measures and any of the cognitive tasks (**Figure 6**). Interestingly, in both the frontal and fusiform regions, MTV and T1 remained stable at varied levels of L2 proficiency (**Figure 2C, F**, **Figure 3C, F**).

**Figure 5.**
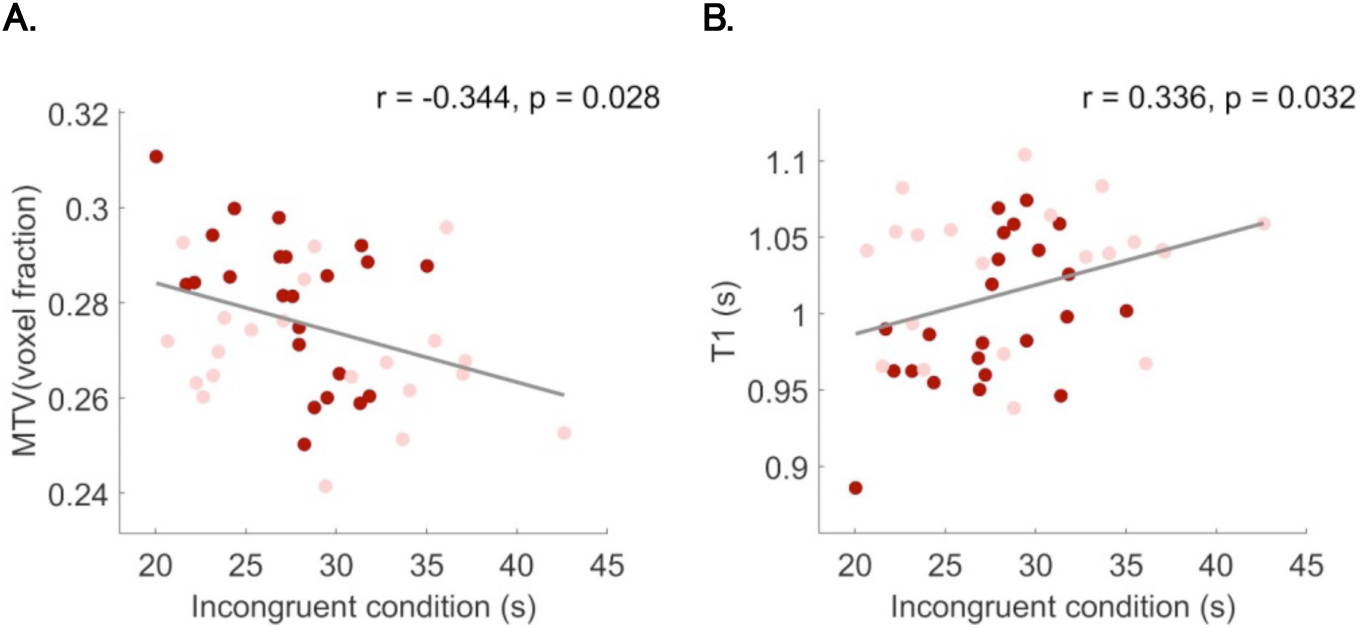
Correlations between total naming time in the Stroop task and qMRI measures in left anterior inferior frontal region. Red dots (n=23) represent early bilinguals and pink dots (n=21) represent late bilinguals. (A) Partial correlation between MTV and total naming time of incongruent English words. (B) Partial correlation between T1 and total naming time of incongruent English words.

**Figure 6.**
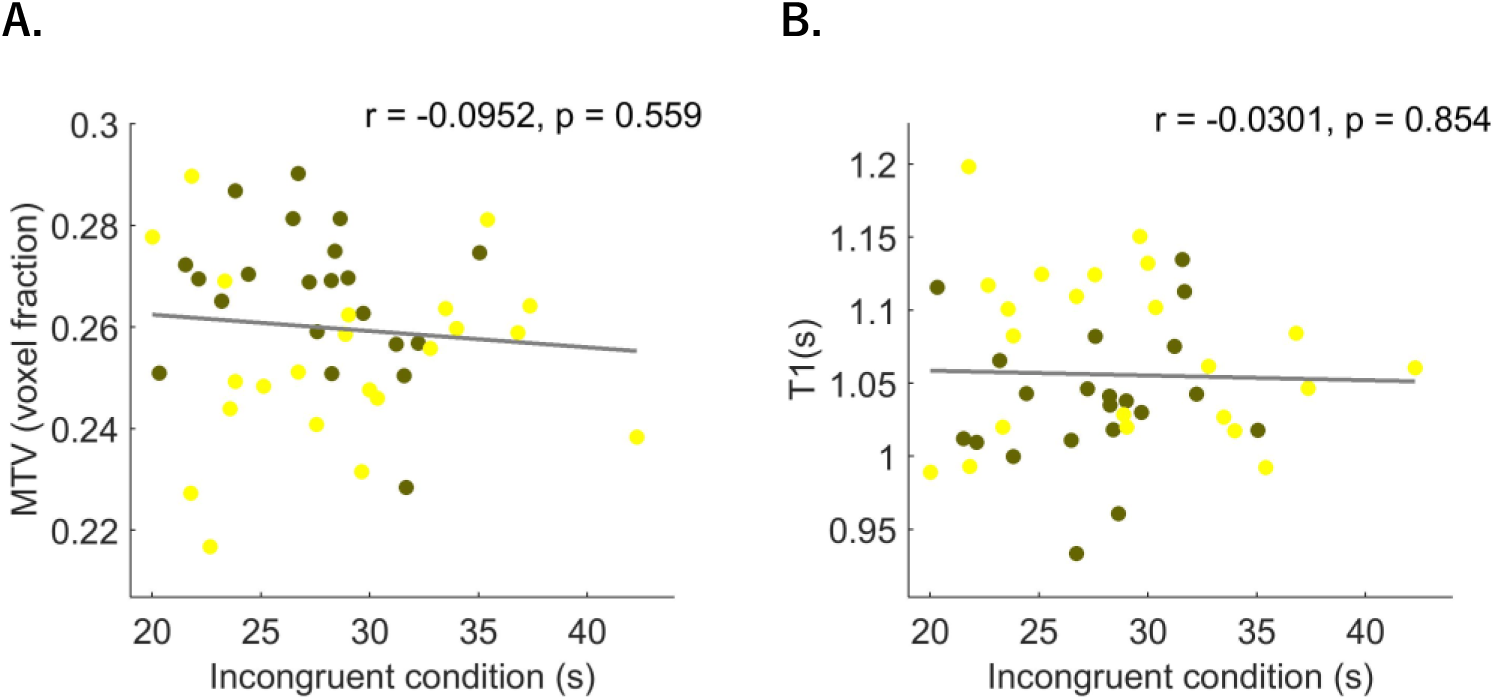
Correlations between total naming time in the Stroop task and qMRI measures in the left middle fusiform region. Olive dots (n=21) represent early bilinguals and yellow dots (n=23) represent late bilinguals. (A) Partial correlation between MTV and total naming time of incongruent English words. (B) Partial correlation between T1 and total naming time of incongruent English words.

## Discussion

Using the qMRI technique, we show in this study that it is possible to identify microstructural plasticity in early bilinguals relative to late bilinguals, especially in key brain regions previously implicated in bilingual language processing such as the anterior inferior frontal and middle fusiform regions. The rapid development of the left inferior frontal region may be related to the inhibition or control of interference processes, as shown in our Stroop task. This finding is highly consistent with previous studies showing that the left inferior frontal cortex is important for bilinguals’ executive functions (Jubin Abutalebi & Green, 2007; Hernandez & Li, 2007; Mårtensson et al., 2012; Stein et al., 2012), a key region also implicated in the AoA effect (Klein et al., 2014; Nichols & Joanisse, 2016). The middle fusiform region is an orthographically sensitive brain region modulated by literacy (Dehaene et al., 2010) and is also related to the competition processes of bilinguals’ two languages (Tan et al., 2011), although we did not find that its microstructural properties (MTV and T1) were correlated with the Stroop effect. Its enhanced development in early bilinguals may have resulted from the need to more efficiently perform orthographic processing in two prints (Liu & Cao, 2016).

Interestingly, our study did not find that language proficiency leads to microstructural neuroplasticity, indicating that L2 proficiency and AoA may have independent effects on neurodevelopment. Previous studies often conflated AoA and L2 proficiency, given the natural confound that early bilinguals also tend to be highly proficient in L2, as compared with late bilinguals (Hernandez & Li, 2007). So far there had been only a few studies that attempted to disentangle the effect of AoA from that of proficiency: Wartenburger et al. showed that these two variables may differentially affect grammatical processing (AoA) versus semantic processing (proficiency) (Wartenburger et al., 2003). In a recent study, Nichols and Joanisse showed with their functional imaging data, consistent with their DTI data, that AoA modulated L2 processing in bilateral IFG inferior frontal gyrus and other regions, whereas proficiency modulated L2 processing in the right cingulate and left parahippocampus, suggesting that these two factors have independent contributions (Nichols & Joanisse, 2016). Our study further confirms that AoA and proficiency can play distinct roles contributing to the microstructure of the bilingual brain.

Our study has demonstrated for the first time that early second language acquisition is associated with enhanced microstructural development in the bilingual brain, and this may provide important evidence for the increased executive functions in early bilinguals compared with late bilinguals or monolinguals (Peristeri et al., 2018). Early learning of a new language seems to lead to microstructural proliferation of the human brain system. One important contribution that this enhanced microstructure makes to better language learning in the early years, according to the ‘sensorimotor integration hypothesis’ (Hernandez et al., 2005; Hernandez & Li, 2007), is that it provides the early learners with an advantage in sensory and motor perception, acquisition, and discrimination, which are critical for acquiring components of a language, including phonetics, orthography, and grammar. The inferior frontal gyrus plays an important role in this process, as it is dedicated to sensory learning, sequence learning, and grammatical and semantic processes (Hagoort, 2005). Previous work has also indicated that the inferior frontal gyrus may form a neural circuity with other regions to accomplish these tasks, including most importantly, the basal ganglia (Ullman, 2001). This neural circuity has been shown to undergo rapid organization in the early years of language development in the context of native language learning (Bates, 2003; Kuhl, 2004), but it has not been examined in greater depth for second language learning. Our findings in the microstructural plasticity sheds light on this issue and provides an avenue for future investigations in this direction.

## Acknowledgments

We thank W. Cui for technical assistance and X.H. Liang for the help in collecting the MRI data. This work was supported by Shenzhen Peacock Team Plan (KQTD2015033016104926), Shenzhen Talent Peacock Plan (827–000115 and 827–000177), Guangdong Pearl River Talents Plan Innovative and Entrepreneurial Team grant (2016ZT06S220), and China’s National Strategic Basic Research Program (“973”) Grant 2012CB720701.

